# Frequency-dependent ecological interactions increase the prevalence, and shape the distribution, of pre-existing drug resistance

**DOI:** 10.1101/2023.03.16.533001

**Authors:** Jeff Maltas, Dagim Shiferaw Tadele, Arda Durmaz, Christopher D. McFarland, Michael Hinczewski, Jacob G. Scott

## Abstract

The evolution of resistance remains one of the primary challenges for modern medicine from infectious diseases to cancers. Many of these resistance-conferring mutations often carry a substantial fitness cost in the absence of treatment. As a result, we would expect these mutants to undergo purifying selection and be rapidly driven to extinction. Nevertheless, pre-existing resistance is frequently observed from drug-resistant malaria to targeted cancer therapies in non-small cell lung cancer (NSCLC) and melanoma. Solutions to this apparent paradox have taken several forms from spatial rescue to simple mutation supply arguments. Recently, in an evolved resistant NSCLC cell line, we found that frequency-dependent ecological interactions between ancestor and resistant mutant ameliorate the cost of resistance in the absence of treatment. Here, we hypothesize that frequency-dependent ecological interactions in general play a major role in the prevalence of pre-existing resistance. We combine numerical simulations with robust analytical approximations to provide a rigorous mathematical framework for studying the effects of frequency-dependent ecological interactions on the evolutionary dynamics of pre-existing resistance. First, we find that ecological interactions significantly expand the parameter regime under which we expect to observe pre-existing resistance. Next, even when positive ecological interactions between mutants and ancestors are rare, these resistant clones provide the primary mode of evolved resistance because even weak positive interaction leads to significantly longer extinction times. We then find that even in the case where mutation supply alone is sufficient to predict pre-existing resistance, frequency-dependent ecological forces still contribute a strong evolutionary pressure that selects for increasingly positive ecological effects (negative frequency-dependent selection). Finally, we genetically engineer several of the most common clinically observed resistance mechanisms to targeted therapies in NSCLC, a treatment notorious for pre-existing resistance. We find that each engineered mutant displays a positive ecological interaction with their ancestor. As a whole, these results suggest that frequency-dependent ecological effects can play a crucial role in shaping the evolutionary dynamics of pre-existing resistance.

## Introduction

The rapid, and often inevitable, evolution of therapy resistance is the primary threat to modern medicine’s successful treatment of cancer, and infectious disease (e.g. bacterial, viral, fungal, and parasitic infections)^1–5^. The story of resistance and treatment failure is strikingly similar across biological kingdoms. A patient is diagnosed and undergoes an initially successful treatment, only for a small resistant subclone of the original disease to relapse, resulting in treatment failure. For decades, the response to this paradigm has been the development of novel, more efficient drugs, targeting orthogonal pathways in hopes of winning the evolutionary arms race. While this response has undeniably resulted in major success stories when considering individual cancers or infections, the overall outlook for drug-resistant disease remains grim^6–9^.

As a result, growing efforts have been made to study these diseases in an *evolutionary* context, whereby scientists seek to understand the ecological and evolutionary forces that seem inevitably to result in the untreatable disease state. Understanding these evolutionary forces that lead to resistance, should allow scientists and physicians to not only design more effective drugs, but perhaps more crucially, design more effect *treatments*. For example, recent work has focused on improving and prolonging the efficacy of our already established drugs via optimal dose scheduling^10–12^, drug combinations^13–17^, understanding spatial dynamics^18–20^, understanding ecological interactions between competing subclones^21–24^, and exploiting collateral sensitivity^25–29^.

In a similar spirit, this work seeks to understand the evolutionary fates of potential resistance-conferring mutations that emerge *before* treatment has occurred. The fraction of these mutants that survive to see treatment are often the primary cause of treatment failure, referred to as “pre-existing resistance”^30–33^. While these resistant populations provide a large fitness advantage once treatment begins, they often carry a significant fitness disadvantage, or fitness cost (*f*_*c*_), in the absence of treatment^34–38^. Nevertheless, resistance-conferring mutants often persist until treatment, at which time their treatment-sensitive ancestors are preferentially killed, resulting in the competitive release and relapse of the resistant population and inevitable treatment failure^39,40^. Understanding how these resistant clones – with a fitness disadvantage – persist in the disease population prior to treatment may allow us to prevent resistance from emerging.

This interest is derived from recent work where we measured the frequency-dependent ecological interaction between an evolved EGFR tyrosine kinase inhibitor (TKI) resistant non-small-cell lung cancer (NSCLC) population and its TKI-sensitive ancestor^40^. The focus of that work was on the ecological interaction under TKI treatment, and the inevitable competitive release. Strikingly, we observed an interaction between the resistant mutant and its ancestor in the absence of any treatment. The resistant population was observed to grow about twenty percent slower than the ancestor when cultured separately, however when the resistant population was co-cultured with a majority ancestor population, that difference in fitness nearly vanished. This observation, referred to as negative frequency-dependent selection (negative because the fitness for the mutant decreases as the mutant frequency *increases*), is a long-studied phenomenon^41–43^, and has been described as the most “intuitively obvious explanation for polymorphisms in nature”^44^. Despite its long history and potential for potent evolutionary effects, frequency-dependent selection remains understudied in the context of drug resistance. This is especially surprising, because a resistant population typically first emerges as a single individual in a predominantly ancestor population, and as a result frequency-dependent ecological interactions have a profound potential to effect the dynamics of a resistant clone (**Fig. 1**).

**Figure 1.**
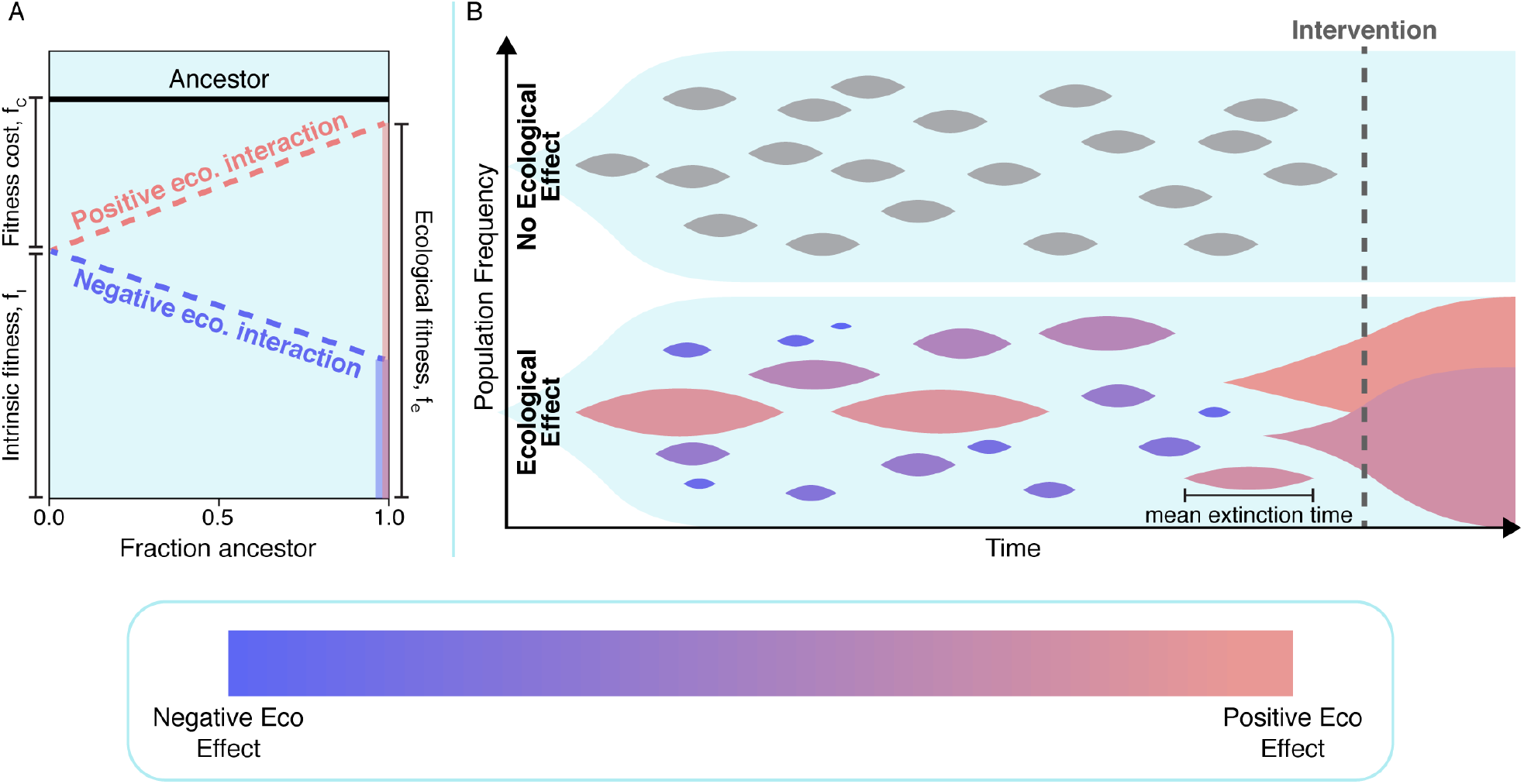
Cartoon abstraction demonstrating how frequency-dependent ecological interactions might increase the likelihood of pre-existing resistance. (A) Cartoon visualization of a typical frequency-dependent growth experiment. The ancestor (black line) is assumed to grow at a constant rate. Two hypothetical resistant mutants are depicted. Both mutants share the same intrinsic fitness and fitness cost, however the positive ecological mutant (red, growth increases as the fraction of ancestor cells increases) has a significantly higher ecological fitness *f*_*e*_ ≈ 1) than the negative ecological mutant (blue, growth decreases as the fraction of ancestor cells increases). (B) Top: Cartoon visualization of an evolving population with no ecological interactions. All mutants are assumed to have a non-insignificant fitness cost, *f*_*c*_, and as a result go extinct. Bottom: The same evolving population, assuming ecological interactions are present. Note that an identical number of mutants emerge, however semi-rare mutants with positive ecological interactions demonstrate an increased time to extinction. As a result, when a drug intervention is administered, pre-existence is much more likely to be present.

In this work, we seek to develop a rigorous theory of pre-treatment evolution that incorporates frequency-dependent ecological interactions between the emerging resistant subclones and the ancestor from which they evolve. Using both a generalized Moran process and Wright-Fisher simulations, we show that mutants with the same intrinsic fitness (monoculture fitness) can have mean extinction times that vary by several orders of magnitude as a function of their ecological fitness (fitness when co-cultured in a predominantly ancestor environment). Next, we calculate the expected number of resistance-conferring mutants in the population as a function of the cost of resistance, as well as the population size, and rate at which resistance-conferring mutations occur. When comparing the result of this calculation both when we assume ecological interactions exist, and when they are forbidden, we identify a wide parameter space where pre-existence is only likely to occur if ecological-interactions are assumed. We then investigate the “many mutant regime” where pre-existence is likely even without ecological interactions, and demonstrate that these ecological interactions would play a prominent role in shaping the distribution of mutants, dramatically increasing the prevalence of mutants with high ecological fitness. Importantly, we show that these ecological effects drive the evolutionary outcomes even when mutants with high ecological fitness are rare. Surprisingly, despite the complexity of the model, we obtain analytical approximations for extinction rates, expected number of resistance-conferring mutants, and the distribution of observed mutants over the full range of ecological fitness. These analytical approximates both support our numerical simulations and allow us to extend our results to population sizes too large to simulate.

Finally, we test our theory experimentally by engineering several of the most common clinically-observed mutations to TKI-therapy in EGFR-driven NSCLC and compete these mutants against the TKI-sensitive ancestor. In all cases we observed an ecological interaction that resulted in mutant ecological fitnesses larger than their intrinsic fitness. Taken together, these theoretical and experimental results argue that frequency-dependent ecological interactions between resistance mutants and their ancestor confer the primary mode by which resistance emerges in modern cancer therapeutics, and potentially all evolutionary diseases.

## Results

### Ecologically-dependent extinction time distributions with a generalized Moran process

We begin by considering a one-step birth-death process^45–47^ with states *s* ∈{0, 1, .., *N*}, where *N* is the total population size, *s* is the mutant population, and *N − s* is the ancestor population. We do not consider mutation and as a result the states *s* = 0 (extinction) and *s* = *N* (fixation) are absorbing. To account for ecological interactions, the mutant’s growth rate is defined to be a function of 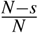, or the fraction of the population that is of the ancestral type, and assumed to be linear. In addition, for simplicity, we define the ancestor’s growth rate to be constant and, without loss of generality, normalized to 1 (see Materials and Methods for full model details). First, we are interested in how the distribution of extinction times differs between recently emerged mutants with identical fitness costs, but distinct ecological interactions (**Fig. 2A, left**). In particular we assume one (neutral) mutant has no ecological interaction with the ancestor, and thus 1− *f*_*c*_ = *f*_*i*_ = *f*_*e*_ (**Fig. 2A, blue**), while the comparative (positive) mutant has an interaction that ameliorates the fitness cost of the mutant at extremely large ancestor fractions, *f*_*e*_ = 1 (**Fig. 2A, red**). Here *f*_*c*_ is the fitness cost or difference between the ancestor’s growth rate and the mutant’s mono-culture growth rate, *f*_*i*_ is the mutant’s intrinsic fitness or mono-culture growth rate, and *f*_*e*_ is the mutant’s ecological fitness or a mutant’s fitness in an otherwise purely ancestor co-culture environment (definitions depicted visually in **Fig. 2A, left**). In population genetics literature this positive interaction is known as negative frequency-dependent selection and fitness is typically plotted as a function of the mutant’s frequency rather than the ancestor. Throughout this work we have chosen to plot fitness relative to the ancestor’s frequency in order to more intuitively connect a positive ecological interaction with a positive slope in frequency-dependent growth plots. In the case of a mutant with a positive ecological interaction, we see that the extinction time distribution is heavily right-skewed in comparison to a neutral ecological effect. As a result, if these two mutants were equally likely to emerge in a population, we would expect to observe a mutant with a positive ecological interaction significantly more often than an equivalent mutant with a neutral ecological interaction. However, ecological interactions are not always positive. Repeating this process in comparing a neutral mutant with a mutant that has a negative ecological interaction with the ancestor reveals a distinct shift to shorter extinction times (**Fig. S1**).

**Figure 2.**
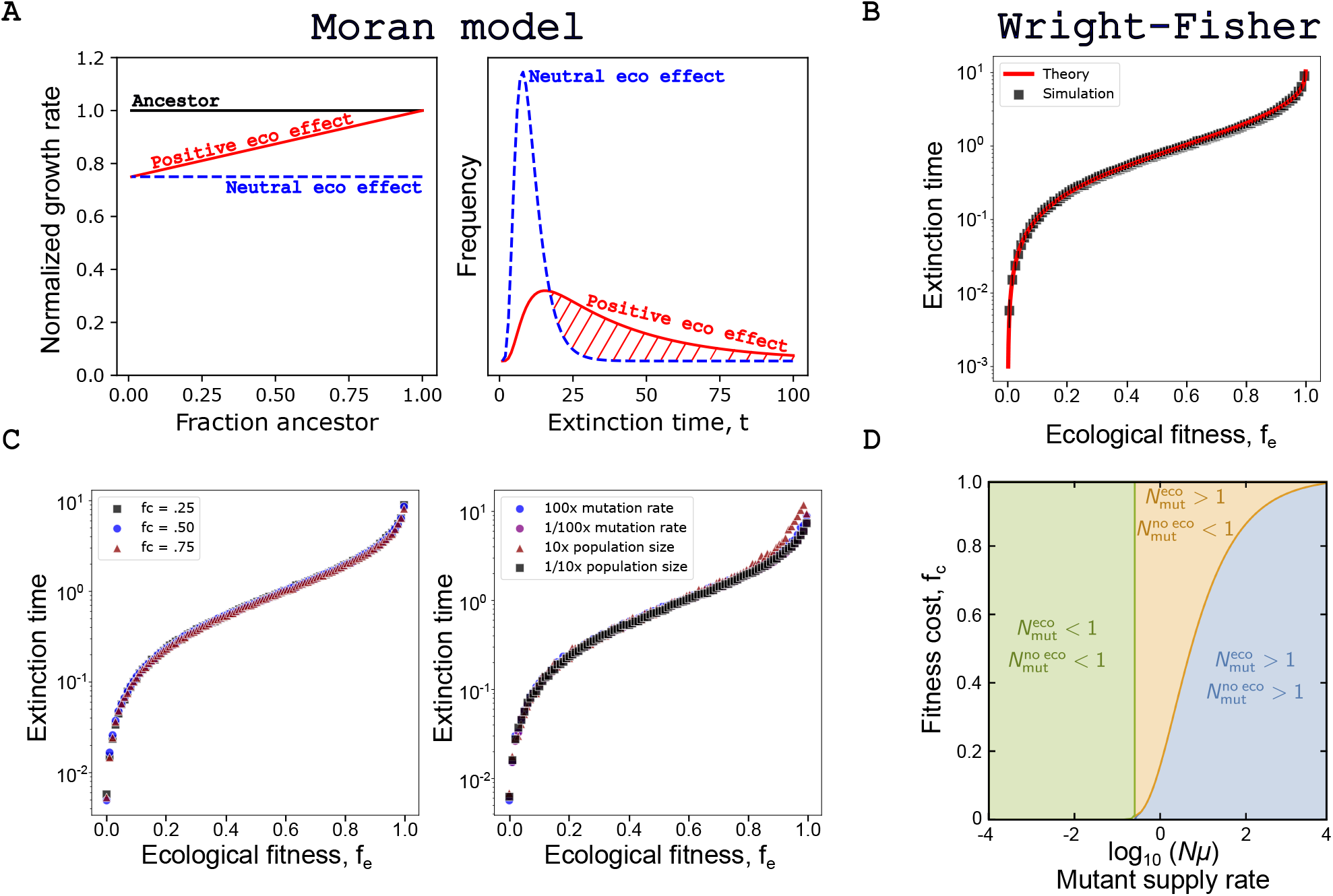
Analytical approximations and simulations predict that extinction times depend on ecological interactions. (A) Closed form extinction time distributions are calculated and visualized for a generalized Moran process (*N* = 100, *f*_*c*_ = 0.25). The red distribution results from a mutant with a positive ecological interaction with the ancestor (*f*_*e*_ = 1.0), while the blue population has no ecological interaction with the ancestor (*f*_*e*_ = 1 − *f*_*c*_ = 0.75). (B) Wright-Fisher simulations are used to numerically calculate the mean extinction time as a function of *f*_*e*_ (*N* = 10000, *µ* = 10^−6^, 500 generations, *f*_*i*_ is drawn uniformly in [0,1 − *f*_*c*_], *f*_*i*_ is drawn uniformly in [0, 1]). (C) Wright-Fisher simulations are repeated for varying values of *f*_*c*_, *µ*, and *N* to confirm theoretical prediction that the extinction time distribution depends only on *f*_*e*_. (D) Phase diagram depicting the three regimes of pre-existing resistance.

### Extinction times depend on ecological interactions in a Wright-Fisher model

While formulating our system as a generalized Moran process allows for convenient closed-form solutions to quantities of interest such as extinction time distributions, this representation becomes computationally expensive as the population size approaches increasingly realistic values. In addition, we have completely ignored mutation, as well as more realistic conditions where many mutants are competing within an evolving population. As such, we switch to a Wright-Fisher formulation of our system^48–50^. In the Wright-Fisher model, populations are still constant in population size *N*, however each individual of the population is replaced every generation with offspring inheriting the parent’s genotype with probability proportional to the parent’s fitness. (Note that in many cases systems with varying populations can be approximately mapped onto a Wright-Fisher model that describes similar evolutionary dynamics, in which case *N* is interpreted as an effective population size^51^.) In addition, individuals acquire mutations with some probability *µ* and we assume mutant populations are sufficiently small that we can ignore both ecological and genetic mutant-mutant interactions. Still, several characteristics from the generalized Moran process remain. Namely, the ancestor’s growth is defined to be constant and normalized to 1, and the mutant growth rate is assumed to vary linearly between *f*_*i*_ and *f*_*e*_ (as a result, a mutant’s growth is fully characterized by these two fitness values along with the fraction of the population that is ancestor).

Each simulation begins with an exclusively ancestor population and with each generation cells mutate with probability *µ*. Each mutant that arises has an intrinsic fitness drawn with uniform probability in [0,1 − *f*_*c*_] and a corresponding ecological fitness drawn with uniform probability in [0, 1]. Each Wright-Fisher ‘generation’ consists of a mutation step, followed by an offspring/selection step. For each mutant that emerges we record its intrinsic and ecological fitness values and track its evolutionary trajectory, and thus extinction time (*τ*). A mutant that emerges but does not survive the subsequent selection step is defined to survive 0 generations. Employing this model we find that the mean extinction time varies nearly five orders of magnitude between the most positive (≈10 generations) and deleterious (≈0.001 generations) ecological interactions (**Fig. 2B**). In order to develop a more rigorous understanding of the evolutionary dynamics, we sought an analytical approximation for the extinction time of a mutant population size, or fitness cost. This finding. Strikingly, we find a robust approximation across the whole range of *f*_*e*_:

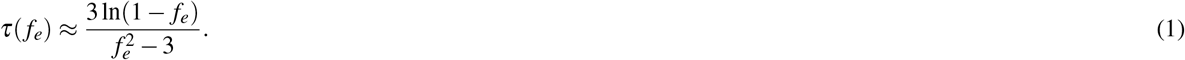

Despite its simple form, this approximation agrees with simulation results with a typical error of 5% (**Fig. 2B**, full derivation and details found in the SI). Interestingly, the approximation is only a function of ecological fitness, and not mutation rate (assuming *µ* ≪ 1), population size, or fitness cost. This finding is supported by our simulation results (**Fig. 2C**). The dependence of *τ* solely on *f*_*e*_ in Eq. (1) is due to two factors: i) the small total proportion of mutants in the population, which means the fitness of a mutant is approximately *f*_*e*_; ii) at each time step (new generation) the chances of the mutants achieving a population comparable to *N* are vanishingly small, so the extinction probability becomes approximately independent of *N*.

### Ecological interactions can increase the probability of pre-existing resistance

Next we consider the model’s implications for pre-existing resistance. Specifically, we are interested in quantifying the expected number of mutants in an evolving population. While it might be tempting to quickly conclude that including ecological interactions will necessarily increase the probability of pre-existing resistance because positive interactions will lead to longer extinction times, it is important to note that mutants with a high intrinsic fitness are more likely to acquire a relatively deleterious ecological fitness, than one that is beneficial. As such, a careful mathematical treatment is required. When the expected number of mutants in a population is low (*N*_mut_ ≪ 1), potential resistance-conferring mutations are unlikely to be present at time of treatment. Contrarily, when the expected number of mutants is greater than 1, we expect treatment threatening resistance to be present when a drug is administered. We begin adapting our analytical model to calculate the mean number of mutants (see SI for full derivation and details). To begin we consider the case where no ecological interactions are present (*f*_*i*_ completely describes the growth rate of the mutant). In this case it can be shown that 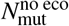, the mean number of mutants ignoring ecological interactions, is:

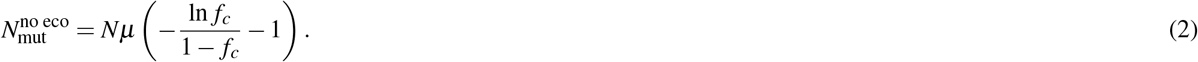

Next, we seek to find an analytical approximation for 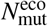, the mean number of mutants assuming ecological interactions exist. In the case of a sufficiently small mutation rate, we can approximate the total mutant fraction as,

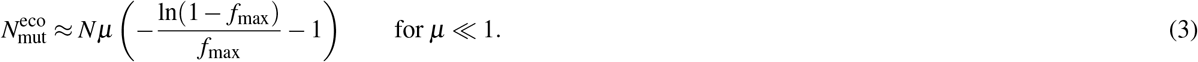

Here, *f*_max_ is the maximum value that *f*_*e*_ can take. While we can set *f*_max_ arbitrarily close to 1, it can never be exactly 1 for a well-defined normalization. Interestingly, for sufficiently small *µ*, the ratio 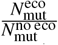 is constant with *Nµ*. While the simplicity of the approximation is appealing, unfortunately it breaks down as *µ* gets large. As a result, a more robust, though significantly more complex, approximation was derived (see SI for full derivation):

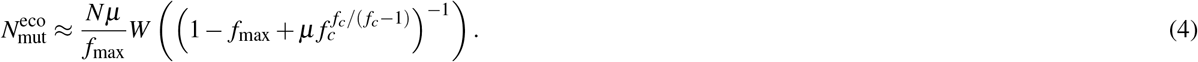

Here *W* (*x*) is the Lambert *W* function, which is the solution *y* of the equation *ye*^*y*^ = *x*. This approximation allows for efficient calculation across several decades of *µ* within 10% of our numerical simulations. Employing these analytical approximations we identify three regimes of interest. The least interesting regime is the small *Nµ* regime (**Fig. 2D, green**).

Here the effective population size is insufficient to maintain a mutant subpopulation regardless of the strength or frequency of ecological interactions. This regime corresponds to extremely rare pre-existing resistance and high likelihood of treatment success.

As *Nµ* gets larger (**Fig. 2D, yellow**) we enter a regime where ecological interactions would suggest pre-existing resistance is likely 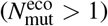, while ignoring ecological interactions would suggest pre-existence is still rare 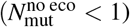. In this regime mutants have yet to become abundant, however, mutants with strong ecological interactions persist sufficiently long to threaten treatment efficacy. Representative simulation trajectories of this “rare mutant regime” are shown in **Fig. 3A**. Without ecological interactions (**Fig. 3A, top panel**) the mutation rate alone is insufficient to maintain a mutant subpopulation capable of threatening future treatment efficacy. However, with the introduction of ecological interactions (**Fig. 3A, bottom panel and Fig. S2**), rare positive ecological mutants climb to significant fractions of the population, and have measurably longer extinction times that may threaten future treatments. As one might intuitively expect, the size of this regime where ecological effects drive pre-existence is heavily dependent on the imposed fitness cost of resistant mutants. We find that the larger the fitness cost imposed by resistance, the larger the comparative increase provided by allowing ecological effects.

**Figure 3.**
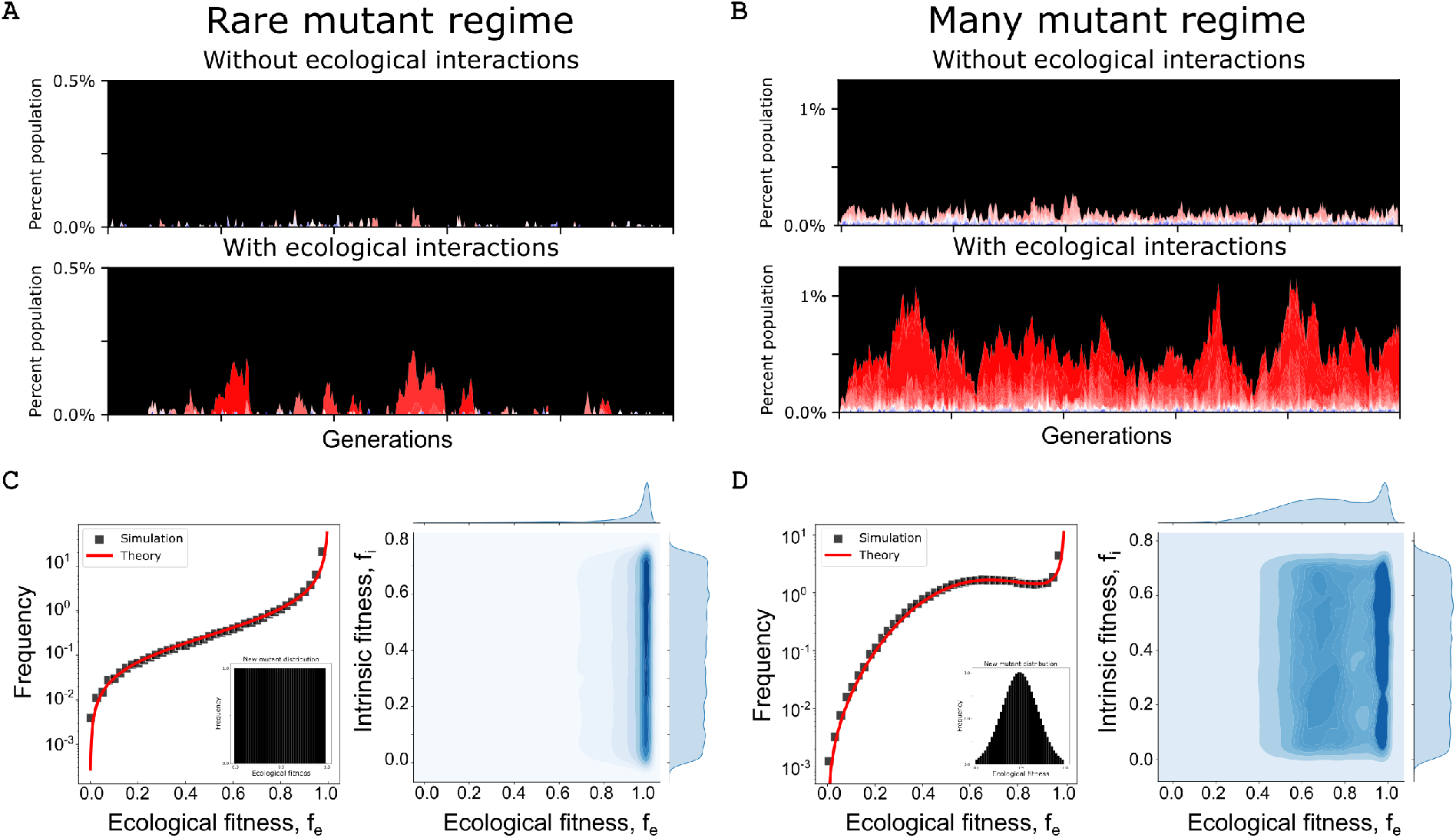
Positive ecological interactions make pre-existence more likely and dominate the stationary distribution of mutants. (A) Representative Wright-Fisher trajectory in the “rare mutant regime”. Black corresponds to the ancestral population. Mutants exist in higher fractions and for longer periods with ecological interactions. Each mutant is colored by its ecological fitness where red represents an *f*_*e*_ value near 1 and blue represents an *f*_*e*_ value near 0. (B) Representative trajectory in the “many mutant regime”. Strong positive ecological interactions dominate the stationary distribution of mutants (visually the mutants appear red, not blue). (C) Left: Stationary distribution of mutant ecological fitnesses when the mutant generating function is uniform across ecological fitness. Right: joint distribution density plot between intrinsic and ecological fitness. (D) Same as C, however the mutant generating function is now Gaussian centered about *f*_*e*_ = 0.5.

### Ecological interactions significantly influence the distribution of mutants

Next, we consider the final regime when *Nµ* is large (**Fig. 2D, blue**). In this regime the mutational supply is sufficiently large to self-sustain a small, resistant subpopulation, regardless of ecological interactions (that is, both 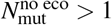 1 and 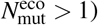. Representative simulation trajectories of this “many mutant regime” are shown in **Fig. 3B**. At first glance one might assume this regime is uninteresting. In both cases mutants are sufficiently common to threaten future treatments, albeit ecological interactions significantly increase the steady-state fraction of resistant mutants. However, the results become more interesting when we consider the shape of the resistant subpopulation distribution. In each trajectory plot, the color is proportional to the mutants ecological fitness with red representing an ecological fitness near 1 and blue representing an ecological fitness near 0. By inspection it is immediately clear that the most positive ecological mutants are over represented in the mutant population, considering they emerge with equal probability. However, we can do better and extract this relationship explicitly from our simulations (**Fig. 3C, left**). We find, similar to the impact of ecological effects on extinction times, that the frequency of a mutant spans multiple orders of magnitude as a mutant’s ecological fitness varies from 0 to 1.

Extending our previous analytical work, it is straightforward to show that the stationary distribution of mutant ecological fitnesses goes as,

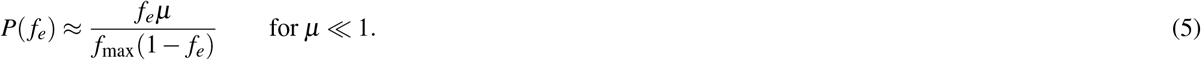

The above approximation works remarkably well despite the simplicity of its form. From this equation we find that the frequency of a mutant is invariant with respect to fitness cost and population size. This is shown explicitly via numerical simulations and visualization of the joint distribution of fitness cost and ecological fitness (**Fig. 2C, right**).

### Non-uniform ecological distributions show similar qualitative results

An important context to keep in mind with the work is that up to this point we have assumed emerging mutants are assigned an ecological fitness with uniform probability in [0, 1]. This assumption was not made for simplicity, but instead out of necessity. While evolutionary biologists have spent significant time both theorizing about, and measuring the distribution of fitness effects (DFE), very little time has been spent quantifying either the frequency or magnitude of ecological effects (which we propose calling the distribution of ecological effects, DEE). As a result, it is difficult to even speculate on what the null model ought to be.

Crucially, the analytical approximations derived herein can be generalized to fit any assumed, or future measured, DEE. While we assumed a uniform distribution, a Gaussian model where the most positive and negative ecological interactions are rare relative to more modest, or non-interacting mutants, may be more accurate. As an example, the general stationary distribution of mutant ecological fitnesses would become,

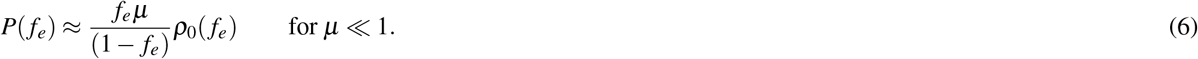

Here, *ρ*_0_(*f*_*e*_) can be any theorized or measured distribution of ecological effects. Eq. (6) can be roughly interpreted as a balancing of two opposing forces to produce a stationary state: the probability ∝ *f*_*e*_*µρ*_0_(*f*_*e*_) that a new mutant with fitness *f*_*e*_ arises and survives is counterbalanced by the probability ∝ (1− *f*_*e*_)*P*(*f*_*e*_) that an existing mutant with fitness *f*_*e*_ disappears. As proof of principle, we numerically simulate the distribution of mutant ecological fitnesses under an assumed Gaussian DEE, and show the above analytical approximation still holds. The results are qualitatively similar to the uniform DEE and, strikingly, despite the rarity of mutants with positive ecological interactions, they still manage to dominate the predicted stationary distribution of mutants (**Fig. 2D**).

### Sufficiently large positive ecological interactions result in a stable fixed point between mutant and ancestor

We now briefly consider the regime wherein the ecological fitness of a mutant can sample values greater than 1. Put another way, when the mutant population emerges, it may emerge into an environment where it out-competes its ancestor. Importantly, all emerging mutants still have a nonzero fitness cost relative to the wild-type and therefore as selection increases their frequency, their relative growth advantage becomes a growth disadvantage, preventing a strong selective sweep. Though our earlier analytical approximations do not apply for *f*_max_ > 1, the numerical simulations are robust in this regime. We find that the majority of stationary distribution mutants are mutants with ecological fitnesses larger than the ancestor, or *f*_*e*_ > 1 (**Fig. 4A**).

**Figure 4.**
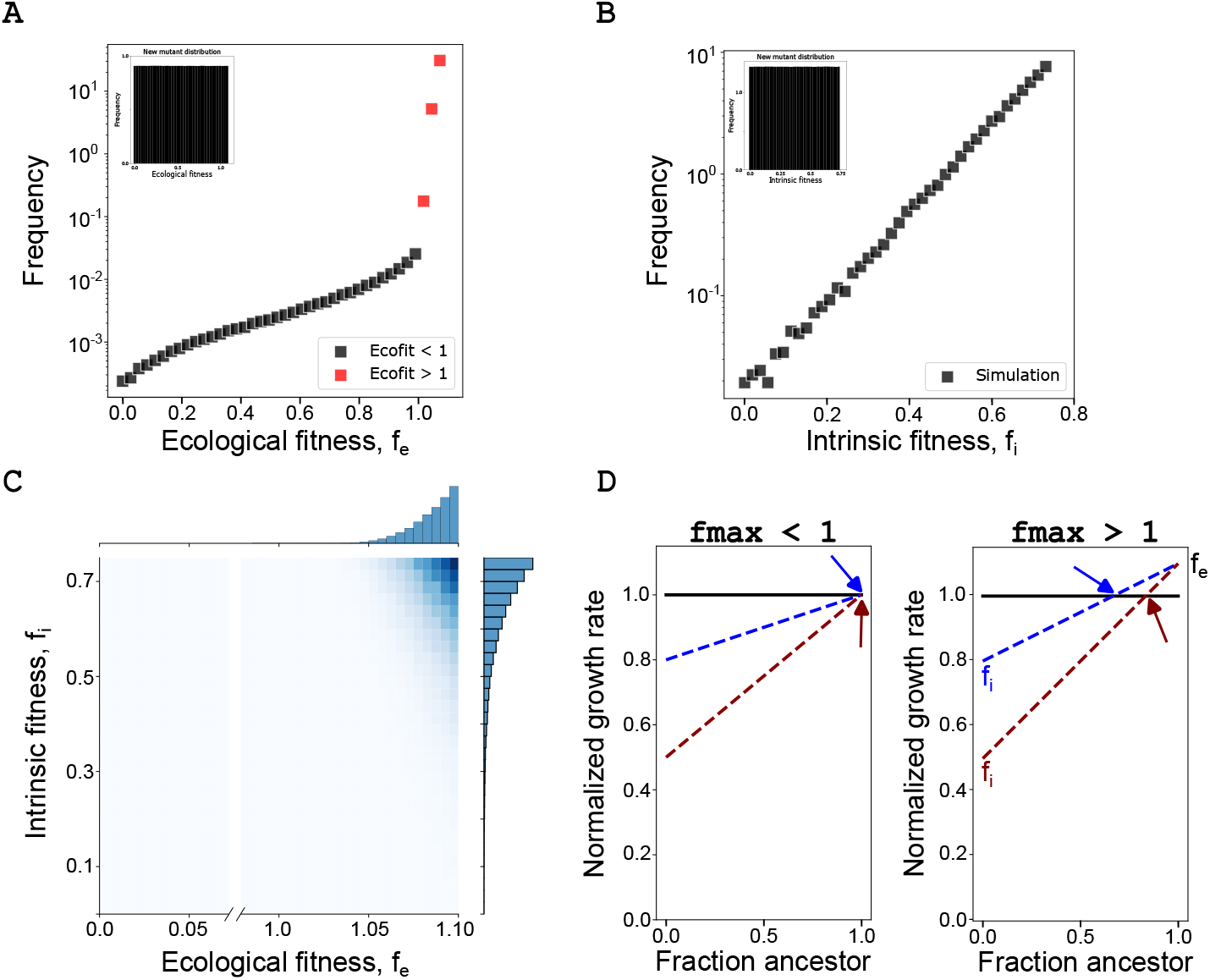
Positive ecological fitnesses above 1 result in a stable fixed point between mutant and ancestor. (A) Stationary distribution of mutant ecological fitnesses when the mutant generating function has uniform probability in [0, 1.10]. (B) Stationary distribution of mutant intrinsic fitnesses. (C) Joint distribution density plot between intrinsic and ecological fitness reveals the size of the fitness cost now has a significant impact on mutant survival. (D) Cartoon illustration of why two mutants with identical values of *f*_*e*_ can result in different extinction times. Colored arrows point to stabled fixed points between mutant and ancestor.

This qualitative change in behavior above *f*_*e*_ = 1 can be explained in evolutionary game theory terms by a switch in the evolutionary game being played. When *f*_*e*_ < 1, the ancestor out-competes the mutant population at all population frequencies. As a result, it is a question of when, not if, the mutant population will be driven to extinction. When *f*_*e*_ > 1, however, the mutant population out-competes the ancestor at high ancestor frequencies, while the ancestor out-competes the mutant at high mutant frequencies (as a result of the mutant fitness cost). This leads to a stable fixed point at some ancestor frequency where the two populations have an equal growth rate. This is particularly worrying in the case of therapy-resistant mutants, because it suggests if such a mutant emerges and survives the initial stochasticity of drift, it will coexist at a sizeable frequency in the population until treatment.

Next, contrary to our previous results, the fitness cost of the mutant plays an important role in determining the stationary distribution of mutants (**Fig. 4B**,**C**). Here we see that only the mutants with the largest positive ecological interactions and smallest fitness costs (intrinsic fitness = 1− fitness cost) are represented at meaningful frequencies. This result can be explained by the qualitative shift in evolutionary game for mutants where *f*_*e*_ > 1. Previously, regardless of the fitness cost, any mutant with *f*_*e*_ ≈ 1 would grow at that ecological fitness, as the mutant population never became a meaningful fraction of the whole population (**Fig. 4D, left**). However, as hinted at in the numerical simulations, *f*_*c*_ and *f*_*e*_ *combine* to determine the stable fixed point between the mutant and the ancestor (**Fig. 4D, right**). As a result, even mutants with *f*_*e*_ < 1 are no longer characterized by their ecological fitness, instead they are characterized by their fitness at the frequency determined by the stable fixed point.

### Clinically observed lung cancer mutations confer positive ecological interactions

Epidermal growth factor (EGFR) tyrosine kinase inhibitors (TKIs) are the first-line treatment for patients diagnosed with advanced non-small cell lung cancer (NSCLC). While the development of targeted TKIs has importantly extended overall survival times, these drugs are rarely curative^39^ and patients often recur with TKI-resistant tumors. As a result, EGFR-mutant NSCLC is an ideal system for studying pre-existing resistance and where we would expect to find positive ecological interactions between mutants and their ancestor.

To test our theory we genetically engineered (see Materials and Methods) three of the most commonly clinically observed resistance mutations found in response to TKIs:^52,53^ BRAF-V600E, KRAS-G12V, and PIK3CA-E545K. Then, using our previously described evolutionary game assay^23,40^, we measured the ecological interaction between each of these mutants and the ancestor PC9 cell line from which they emerged.

Excitingly, we found that each of the three engineered mutants had positive ecological interactions with their ancestor (**Fig. 5, bottom**). In addition, these positive ecological interactions are strikingly similar in both magnitude and shape to the measured ecological interaction between the evolved mutant and its ancestor (**Fig. 5, top**). While we already reported on the ecological interaction between the evolved mutant and its ancestor as it was the motivator of this study^40^, we performed additional sequencing analysis (WXS and RNA-seq) and identified several common clinical mutations present, distinct from the engineered mutations: MET overexpression, CCND1 amplification, and KRASG12D mutation (**Fig. 5, top**). Taken together, the engineered and evolved mutants combine to survey approximately 70% of clinically known resistance mechanisms to TKIs in NSCLC, and in all cases we observed large positive ecological interactions ameliorating a sizeable fitness cost of resistance. Next, as a control, we measured the ecological interaction between the engineered BRAF-V600E and KRAS-G12V cell lines in co-culture and did not observe a positive ecological interaction (**Fig. S3**). This is unsurprising because a key aspect of our model is that positive ecological interactions are enriched during co-evolution and thus should be more likely between ancestor and mutant, not two mutants. These experimental results are harmonious with our theoretical work and strongly support the hypothesis that frequency-dependent ecological interactions can play a critical role in the acquisition of resistance in evolutionary diseases.

**Figure 5.**
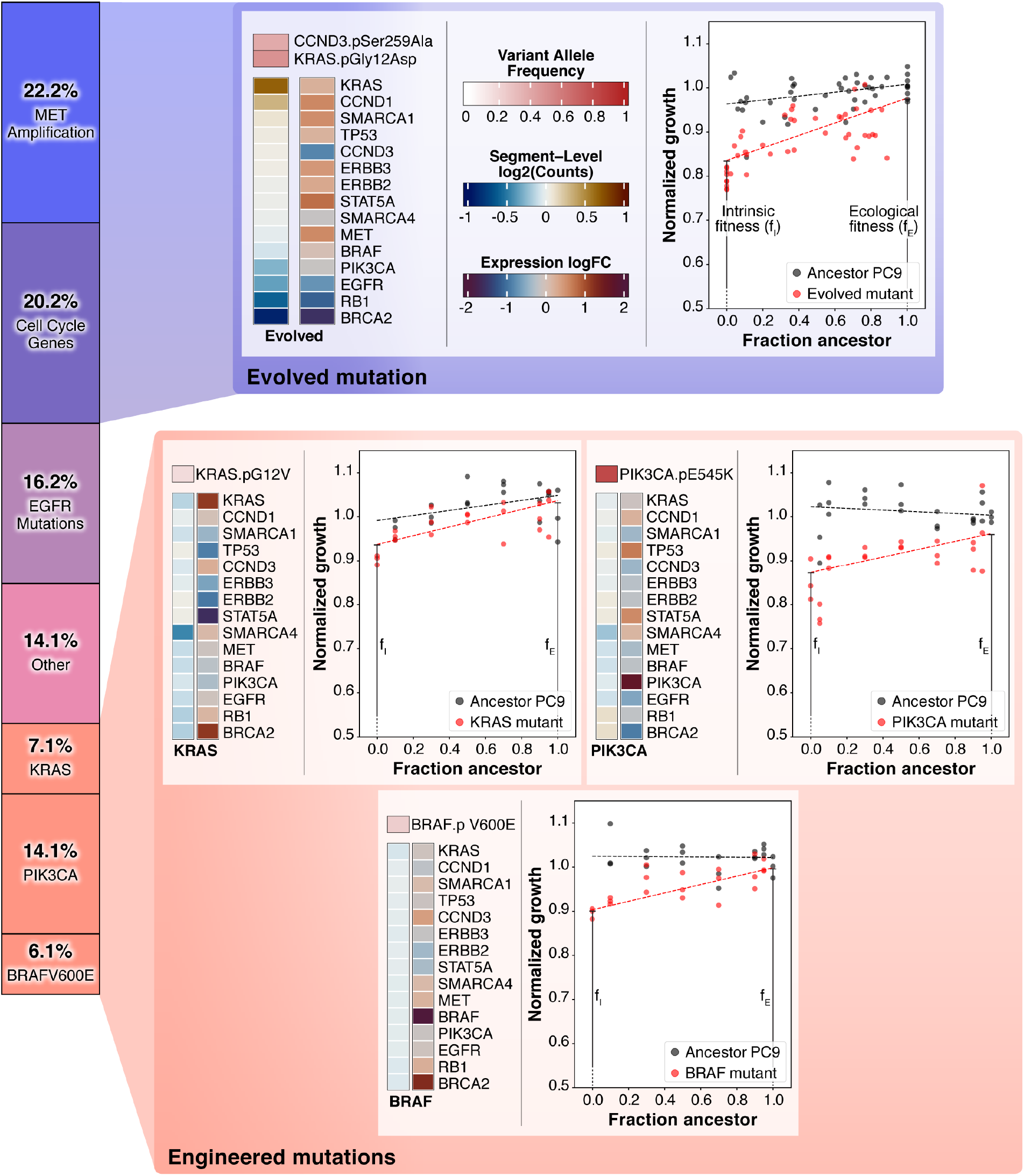
Common clinically observed resistance mutations in NSCLC harbor strong postitive ecological interactions with their ancestor in a model system of pre-existing resistance. Stacked bar chart: Visual representation of the known resistance mechanisms to Osimertinib, a third generation TKI and the current standard of care for EGFR-positive NSCLC. Mutation frequencies and categorical definitions from Leonetti et al^54^. Top: Evolved gefitinib-resistant NSCLC PC9 mutant (previously reported^40^) exhibits a positive ecological interaction with its ancestor. Fresh sequencing analysis identifies clinically observed resistant mutations including: KRASG12D, MET amplification, and CCND1 amplification (cell cycle gene). Bottom: Measured positive ecological interactions between engineered resistant mutants (KRAS-G12V top-left, PIK3CA-E545K top-right, BRAF-V600F (bottom) and their ancestor.

## Discussion

While much work has gone into quantifying clinically problematic resistant bacteria, cancers and viruses, we nearly always characterize these clones in monoculture - entirely outside the eco-evolutionary forces that selected for (or against) them in the first place. In this work we set out to provide the foundation for a rigorous and generalizable mathematical framework that incorporates frequency-dependent ecological interactions and can be used to study their role in pre-existing resistance. This work both compliments and builds off of recent studies from a wide range of disciplines ranging from theoretical population genetics and ecology to clinical trials across several biological kingdoms. We demonstrate that the presence of ecological interactions can significantly increase the probability of pre-existing resistance, in addition to shaping the distribution of mutants likely to be present before treatment. We derive analytical approximations of several quantities of interest including extinction time, mean mutant population numbers, and the underlying distribution of mutants each as a function of ecological fitness. Importantly, these results can easily be generalized to any theorized distribution of ecological effects, or future experimentally measured distribution. As an important example, we show that even when we assume positive ecological interactions are rare, they still end up as a plurality of the stationary mutant frequency distribution. Finally, in an model system for pre-existing resistance, we show common clinically observed mutants harbor positive, frequency-dependent ecological interactions when co-cultured with their ancestor, providing strong evidence for our theory in cancer. In addition, recent exciting work in bacteria provides additional evidence, as frequency-dependent interactions resulted in maintenance of otherwise costly antibiotic resistant populations in *Escherichia coli*^55^ and *Pseudomonas aeruginosa*^56^.

It is also important to address several limitations of our work. As we mentioned earlier, the distribution of ecological effects (DEE) has never been experimentally measured. As a result, assumptions regarding the distributional parameters have to made in order to calculate meaningful quantities of interest. While we did our best to combat this by developing analytical models that are agnostic to this distribution, the quantitative aspect of our results are subject to the specifics of a model. Our hope is that the analytical and numerical results herein, when combined with the promising experimental work in NSCLC, motivate future measurements of the DEE across diverse model systems. Similarly, our own experimental validation is constrained to one subsystem. Our predictions are broad and should apply to many evolving populations where pre-existence is evolved. Therefore it is important that future studies should should aim to test these theories not just in other cancers, but in other organisms from HIV to drug resistant bacteria. It is possible that these principles provide the most explanatory power in cancer and bacteria where it is common to find highly dense heterogeneous populations in contrast to viruses, for example. Finally, while the model aims to generally capture major evolutionary forces that may underlie pre-existing resistance, it is still an abstraction of a much more complex clinical scenario where the immune system, spatial dynamics, and treatment adherence, to name only a few, can play major roles.

## Materials and Methods

### Cell culture

PC-9 are human adenocarcinoma cells derived from undifferentiated lung tissue were obtained from Sigma (Sigma, USA). PC-9 cells were cultured in RPMI-1640 medium supplemented with 10% heat inactivated fetal bovine serum (FBS) and 1% penicillin streptomycin solution at 37°C with humidity containing 5% CO2. Cells were split every four days to maintain optimum confluency of ≈80-90%.

### Engineering of mutant cell lines

To establish PC-9 cells stably expressing target genes, HEK-293T cells were co-transfected using TransIT-Lenti transfection reagent (Mirus, USA), with 500ng psPAX2 (addgene, USA), 100ng PMD2 (addgene, USA) and 500ng of target genes. Viral particles were collected after 48hrs and used to transduce PC-9 cells. Then, to establish ancestor PC-9 cells stably expressing nuclear localized GFP, cells were transduced with pLVX-eGFP-Hygro (Vectorbuild, USA). In addition, to establish query cells expressing fluorescently labeled PC-9 cells with a gene of interest cells were co-transduced with pLVX-mCherry-Hygro or pLVX-mCherry-Puro and each of pLVX-PIK3CA-E545K-Bsd (Vectorbuild, USA). Next, 72hrs after transduction, cells were selected with 200*µ*g/ml hygromycin, 5*µ*g/ml puromycin and 5*µ*g/ml blasticidin.

### Drug sensitivity assay

Cells were harvested at 70-80% confluence, stained with trypan blue (Corning, USA), and counted with a TC20 Automated Cell Counter (Bio-Rad, USA). Luminescent based cell viability assays using CellTiter-Glo (CTG) reagent (Promega, USA) were performed in 96 well plate (Corning, USA). A total number of 3, 000 cells were plated in 90µL of complete medium per well in three replicate per drug concentration with Multidrop reagent dispenser (Thermo Fishers, USA). After 3hrs of incubation, 10µL of gefitinib, osimertinib and erlotinib (Cayman, USA) diluted in complete RPMI-1640 medium were added to the cells. Compounds were tested in a threefold dilution in a range of 0 − 1.8*µ*M, 0 − 3*µ*M and 0 − 10*µ*M for gefitinib, osimertinib and erlotinib respectively. After 72hrs of incubation, 25µL CTG reagent was add to the cells; incubated for 10 minutes at room temperature and luminescence was measured.

### Game assay

PC-9 mutants stably expressing nuclear localized fluorescent signal ancestor PC-9 stably expressing nuclear localized GFP were co-cultured at different initial proportion of ancestor cells at a density of 1, 500 cells in 90*µ*L of fresh medium. After 3hrs of incubation, 10*µ*L of DMSO diluted in complete RPMI-1640 medium (final DMSO concentration of 0.1% v/v) were added to the cells in three replicates per initial proportion. Then time-lapse microscopy images were obtained for GFP and mCherry using BioSpa automated incubator (BioTek, USA) every 4 hours over the course of 96 hrs. Then, images were processed with the open-source software CellProfiler^57^. Images were background subtracted, converted to 8-bit, contrast enhanced, and thresholded, then raw cell numbers were extracted.

### Generalized Moran model details

In this work we consider a well known generalized Moran process, a model previously used to study frequency dependent evolutionary dynamics^45–47^. Briefly, we consider a one-step birth-death process with states *s* ∈{0, 1, .., *N*} and characterized by birth and death rates *b*_*i*_ and *d*_*i*_ with *I* ∈ {1, 1, .., *N* − 1}. As a result this model describes a fixed population size *N*, with *s* resistant mutants and *N* − *s* ancestor population. We forgo mutation rate and as a consequence the states *s* = 0 and *s* = *N* are absorbing. We consider an evolutionary game with a 2*x*2 payoff matrix such that:

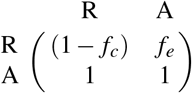

As a result, we can write the expected payoffs in a population of *s* mutants and *N* − *s* ancestor individuals as:

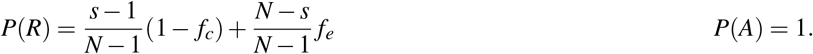

### RNA-Seq

Paired-end reads are preprocessed using *fastp* to trim and quality filter the reads. Following the filtering, reads are aligned to GRCh38 reference genome via *STAR* aligner. Read quantification is done using *Salmon* on the extracted transcriptome locations from spliced STAR alignment. Gene-level abundance are aggregated from bootstrapped transcript abundances using R package *tximport*. The *Arriba* tool is coupled with spliced alignments for fusion-transcript detection as well. Pathway level expression activities are quantified using R package *GSVA* and *msigdbR* for the Hallmark Pathways. The R package *ComplexHeatmap* was used to generate heatmaps.

### Whole exome sequencing

Paired-end whole-exome reads of ancestor and parental lines were preprocessed using *fastp* similar to RNA-Seq. Alignment to GATK (GATK best practices bucket) version of GRCh38 reference is done using *bwa-mem* aligner. Following the alignment, variant calling pipeline according to the GATK workflow including duplicate marking and variant calling via HaplotypeCaller was conducted. Variants passing filtering based on hard-filtering are further annotated using Variant Effect Predictor (VEP) tool. Exome alignments are further input to *CNVkit* for copy-number alterations. Using a flat-reference for bias correction log2 scaled abundances are generated for ancestor and resistant strains. Copy number segments are captured using circual binary segmentation and assigned to genes mapping to the segment.

### Code availability

Code used in this study will be made openly available on GitHub.

## Acknowledgments

This work was made possible by the National Institute of Health T32CA094186 (JAM), 5R37CA244613-03 (JGS), 5T32GM007250-46 (JGS), The Research Council of Norway 325628/IAR (DST), and American Cancer Society RSG-20-096-01 (JGS).

## Supplemental Material

### Materials and Methods

#### Generalized Moran model details

In this work we consider a well known generalized Moran process, a model previously used to study frequency dependent evolutionary dynamics^45–47^. Briefly, we consider a one-step birth-death process with states *s* ∈ {0, 1, .., *N*} and characterized by birth and death rates *b*_*i*_ and *d*_*i*_ with *i* ∈ {1, 1, .., *N* − 1}. As a result this model describes a fixed population size N, with *s* resistant mutants and *N* − *s* ancestor population. We forgo mutation rate and as a consequence the states *s* = 0 and *s* = *N* are absorbing. We consider an evolutionary game with a 2*x*2 payoff matrix such that:

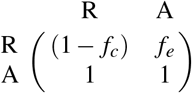

As a result, we can write the expected payoffs in a population of s mutants and N-s ancestor individuals as:

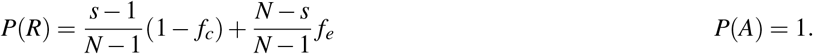

### Simulations

Here we provide additional complimentary results from our numerical simulations in the main paper.

#### Additional generalized Moran process results

Similar to the main text figure, we compare the extinction time distributions of two emerging mutants in an initially ancestor population. Here we compare a neutral mutant with a negative mutant.

While the effect is smaller that the comparison between positive and neutral mutants in the main text, we find similar qualitative trends in that extinction time distribution for neutral mutants is shifted to longer extinction times when compared to the negative mutant (**Fig. S1**).

#### Visualization of sample evolutionary trajectories on a log-axis

Similar to the main text figure, we visualize representative evolutionary trajectories under the rare and many mutant regimes. This time, we the y-axis is on a log scale, which better highlights the presence of low-frequency deleterious mutants in the trajectories (**Fig. S2**).

#### Mutant-mutant game assay control experiment

Each of the three engineered cell lines (BRAF, KRAS, PIK3CA) and the evolved gefitinib-resistant cell line exhibited a positive ecological interaction when co-cultured with the gefitinib-sensitive ancestor PC9 cell line. Our analytical approximations and simulations suggest this is because of the strong selective advantage the positive ecological interactions convey to the resistant mutants. That is, if the pool of drug-resistant mutants contains mutants with strong positive ecological interactions, they will be selected for. However, this does not suggest that these mutants will exhibit ecological interactions with one another, as the selective pressure to confer such an interaction is missing. We hypothesized that the engineered BRAF and KRAS mutants would not exhibit the same positive ecological interaction we observed between each mutant and their ancestor. This hypothesis turned out to be right in this instance (**Fig. S3**). In addition, this experiment serves as a technical control experiment for the game assay, as it reinforces the observed ecological interactions are not a technical artifact (such as one that may arise from a finite error rate in counting of fluorescent cells).

**Figure S1.**
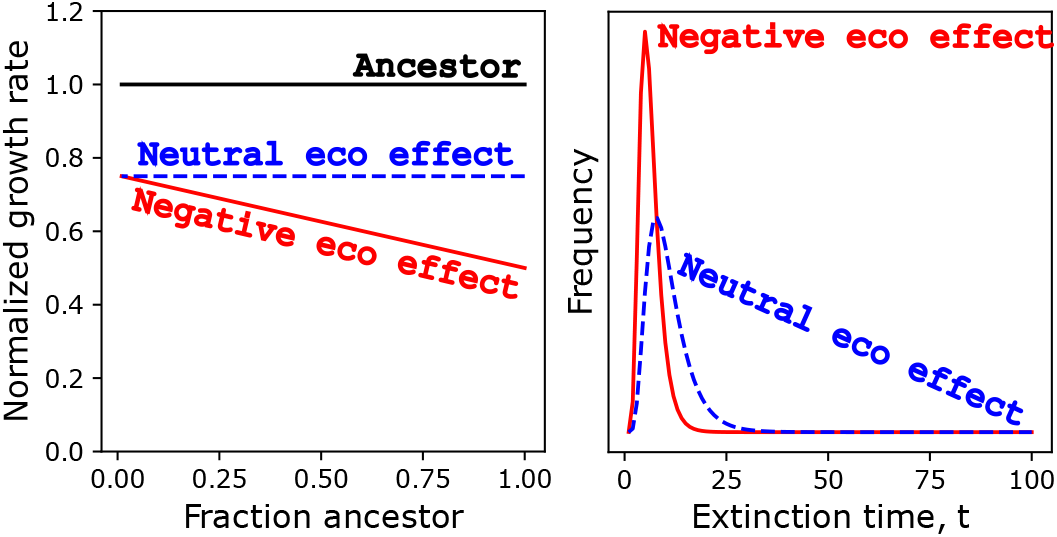
Similar qualitative trends exist when comparing neutral and negative mutants. Closed form extinction time distributions are calculated and visualized for a generalized Moran process (N=100, *f*_*c*_ = 0.25). The red distribution results from a mutant with a negative ecological interaction with the ancestor (*f*_*e*_ = 0.5), while the blue population has no ecological interaction with the ancestor (*f*_*e*_ = 1 − *f c* = 0.75).

**Figure S2.**
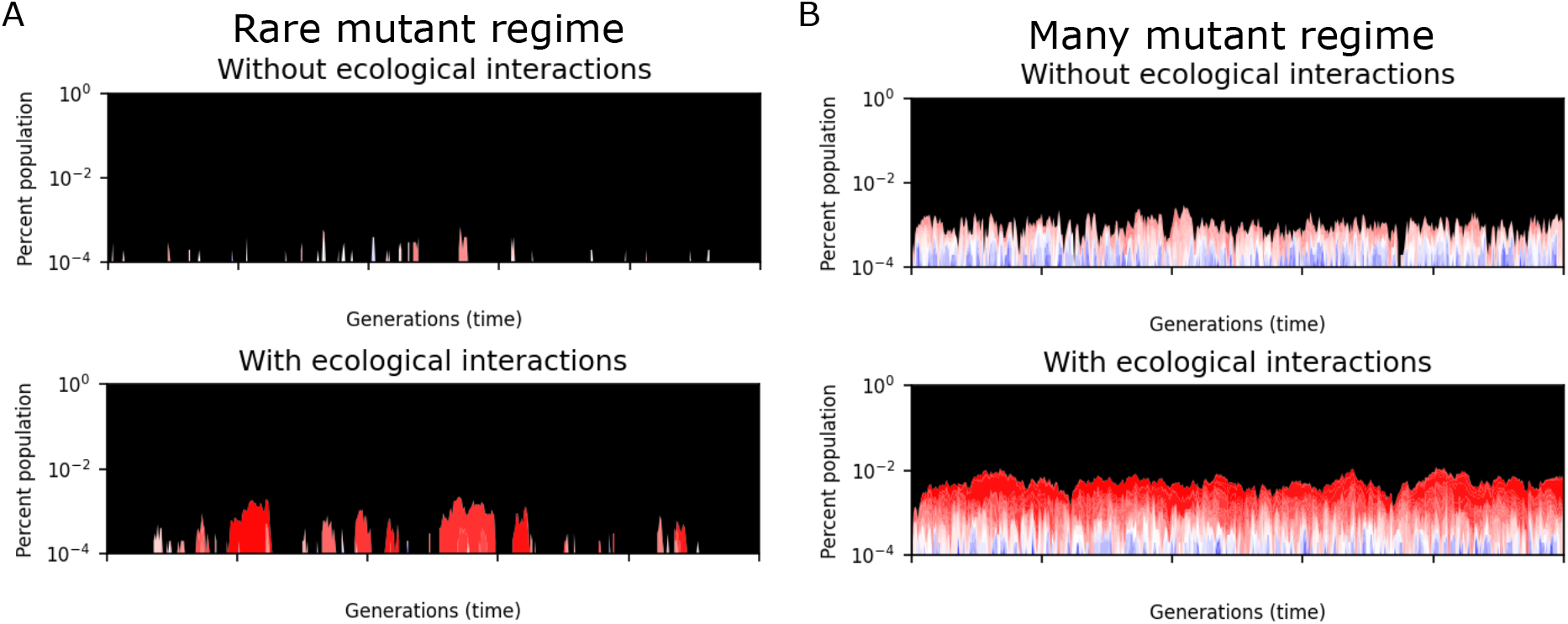
Positive ecological interactions make pre-existence more likely and dominate the stationary distribution of mutants (log-scale). (A) Representative Wright-Fisher trajectory in the “rare mutant regime”. Black corresponds to the ancestral population. Mutants exist in higher fractions and for longer periods with ecological interactions. Each mutant is colored by its ecological fitness where red represents an *f*_*e*_ value near 1 and blue represents an *f*_*e*_ value near 0. (B) Representative trajectory in the “many mutant regime”. Strong positive ecological interactions dominate the stationary distribution of mutants (visually the mutants appear red, not blue).

#### Analytical theory

The following sections describe the analytical theory supporting the numerical simulations in the main text. Important formulas that are used to fit the simulation results are highlighted by boxes.

##### Stationary distribution and total number of mutants

We seek to derive an approximate analytical expression for the stationary probability density of mutants *P*(*f*) in the Wright-Fisher simulations described in the main text. We assume ecological interactions are present, and that sufficient time has passed for a stationary state to be reached. *P*(*f*)*d f* is defined as the fraction of the total population that consists of mutants with instantaneous fitnesses between *f* and *f* + *d f*. Integrating this distribution gives the mean total number of mutants 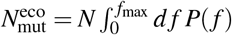, where *N* is the fixed total population size. For the theoretical calculations we consider an upper bound on the mutant fitness *f*_max_ < 1 to allow for a well-defined normalization (as explained in more detail below), though we can set *f*_max_ arbitrarily close to 1 in order to fit the numerical results.

Every generation of the model consists of a mutation step followed by selection. Let us consider first the mutation part. If *P*(*f*) is the current distribution, mutation modifies it to a new distribution,

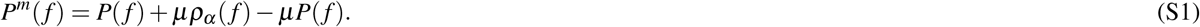

Here *µ* is the mutation probability for a single cell in one generation, and *ρ*_*α*_ (*f*) is the probability density of a new mutant having fitness *f*, given a total fraction of mutants 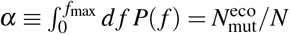. The second and third terms on the right-hand side in **Eq**. (S1) are respectively the gain and loss due to new mutations.

**Figure S3.**
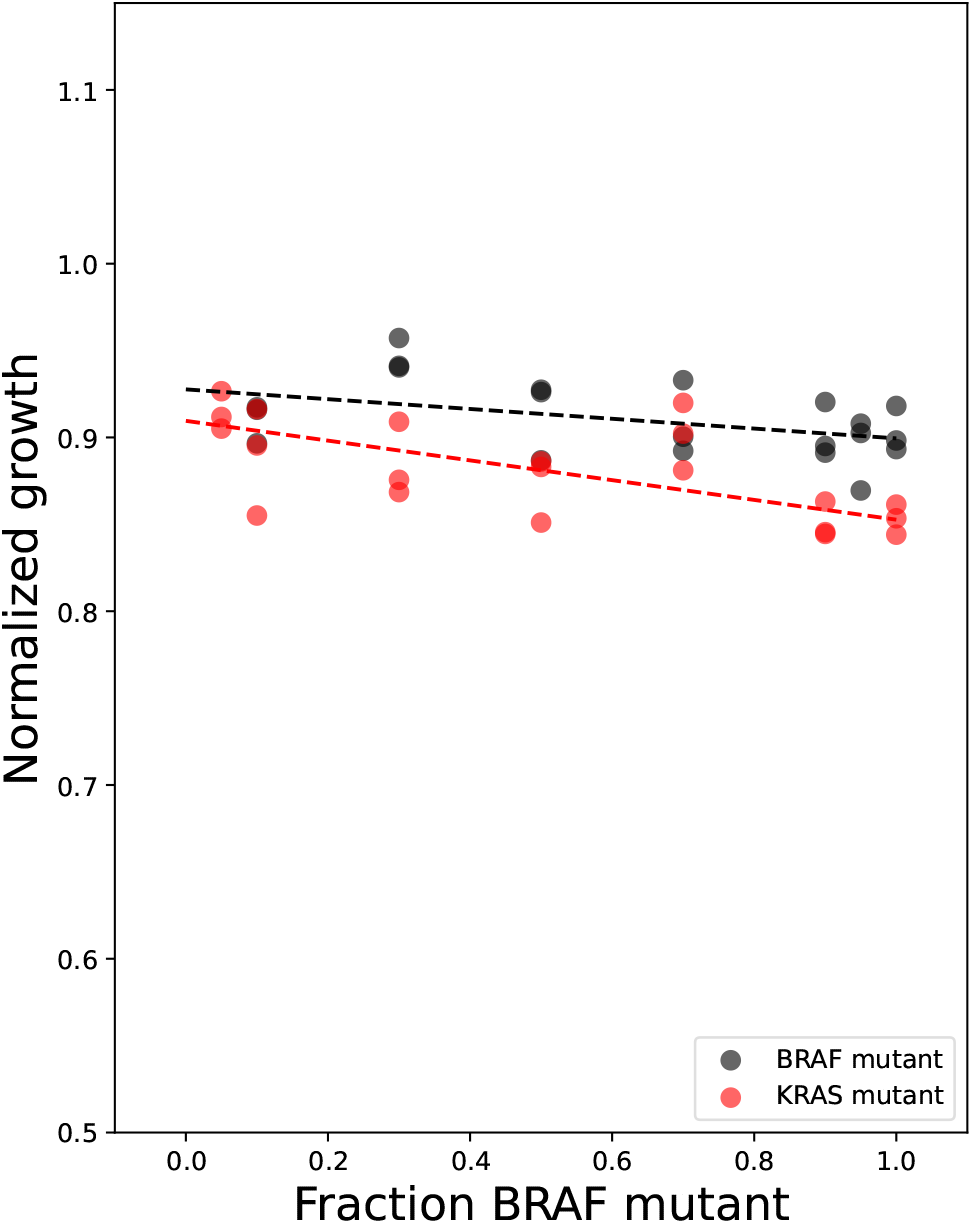
Engineered BRAF and KRAS cell lines do not exhibit a positive ecological interaction when co-cultured. Frequency-dependent growth rate measurements between an engineered BRAF cell line (black) and an engineered KRAS cell line (red).

The dependence of *ρ*_*α*_ (*f*) on *α* reflects the role of ecological interactions. In the simplest linear model, a new mutant is assigned a fitness function *f* (*α*) = *α f*_*i*_ + (1 − *α*) *f*_*e*_, where *f*_*i*_ (the intrinsic fitness) is randomly drawn from a uniform distribution between 0 and 1 − *f*_*c*_, and *f*_*e*_ (the ecological fitness) is randomly drawn from a uniform distribution between 0 and *f*_max_. Here *f*_*c*_, where 0 < *f*_*c*_ < 1, is the cost associated with intrinsic fitnesses, with 1− *f*_*c*_ < *f*_max_. For this definition of *f* (*α*), the distribution *ρ*_*α*_ (*f*) is given by

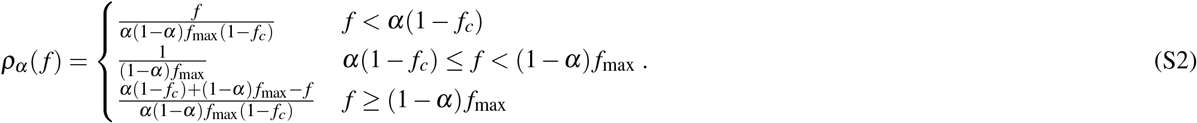

For 0 < *α* < 1 the distribution *ρ*_*α*_ (*f*) has a trapezoidal shape, rising linearly from zero for small *f*, then plateauing in the middle region, before decreasing linearly to zero at *f*_max_. In the two limits *α* = 0 and *α* = 1 it reverts to a uniform distribution between 0 and *f*_max_ (for *α* = 0) or between 0 and 1 − *f*_*c*_ (for *α* = 1).

The second part of the dynamics is the selection step, which makes a further modification of the mutant distribution, yielding

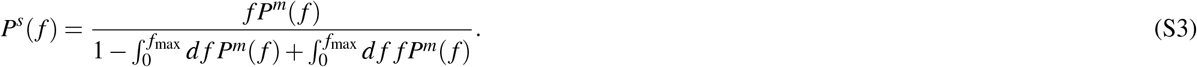

The above form reflects the fact that the ancestors after the mutation step, comprising a fraction 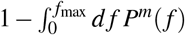 of the total, have fitness 1, and mutants with fitness *f* have a chance of surviving into the next generation proportional to *f*. In order for the system to be in a stationary state, the distribution *P*^*s*^(*f*) in the next generation must end up being the same as the starting distribution *P*(*f*) in the current generation. Using **Eqs**. (S1)**-**(S3), we can express the condition *P*^*s*^(*f*) = *P*(*f*) compactly as

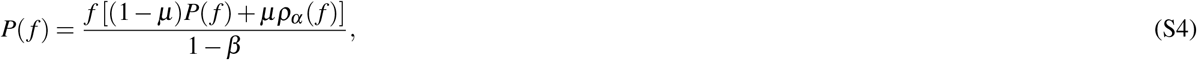

wher 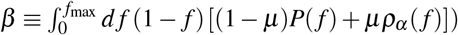 **Eq**. (S4) can be solved for *P*(*f*),

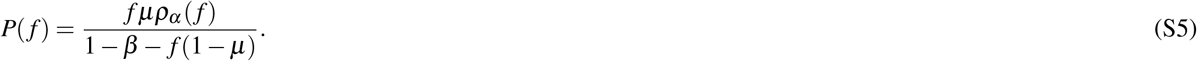

Note that both *α* and *β* on the right-hand side of **Eq**. (S5) depend implicitly on *P*(*f*). In order for the solution to be self-consistent, we plug **Eq**. (S5) into the definitions of *α* and *β*, which leads to a closed system of equations for these two quantities:

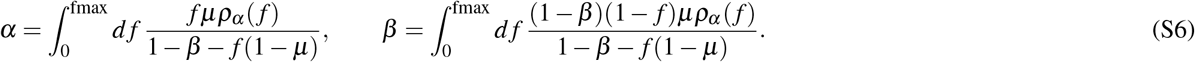

The equation for *β* is satisfied exactly when *β* = *µ*, using the fact that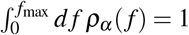. Plugging *β* = *µ* into the *α* equation, we can carry out the integral analytically, leading to the following relation:

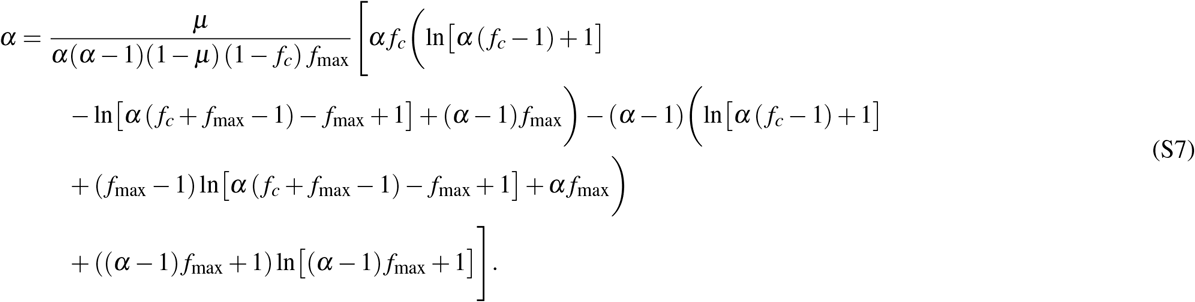

There is no explicit analytical solution for *α* from the above equation, but there are ways to derive approximate solutions that work in different limits. We consider two such limits in turn.

##### Small mutant fractions (*µ* ≪ 1)

When *µ* ≪ 1, the mean number of mutants in the population becomes small, and their fraction *α*, given by the solution of **Eq**. (S7), scales like *α* ∝ *µ*. In this limit **Eq**. (S7) gives us 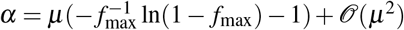 so we get an approximate expression for the total mutant fraction (main text **Eq**. (3)):

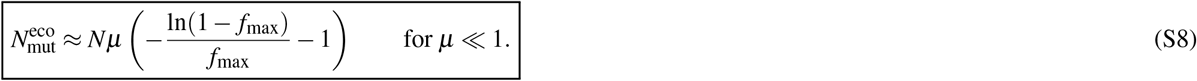

To leading order in *µ*, we can approximate the stationary distribution of mutant fitnesses in **Eq**. (S5) as (main text Eq. (5)):

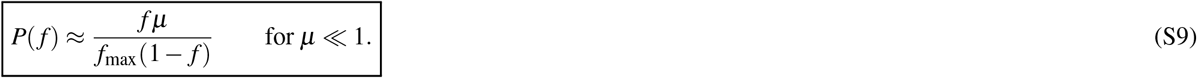

Note that since *α* ≪ 1, *ρ*_*α*_ (*f*) ≈ *ρ*_0_(*f*), and the distribution of instantaneous fitnesses *f* in **Eq**. (S9) is approximately also the distribution of ecological fitnesses *f*_*e*_. In order for **Eq**. (S9) to be normalizable, and hence its integral giving **Eq**. (S8) to be well-defined, we need *f*_max_ strictly smaller than 1. In the numerical simulations only a finite number of mutants are sampled overall during the course of the evolutionary trajectories, and an effective value of *f*_max_ ≈ 0.995 was found to provide good fits between the theory and simulation results.

##### Large mutant fractions (*µ* ≲ 1)

We would like to extend the results above to larger values of *µ* and *α*, to cover simulation cases where the mutant fractions *α* are on the order of 10% of the total. We note that as *α* gets larger, the distribution of new mutant fitnesses *ρ*_*α*_ (*f*) in **Eq**. (S2) gets increasingly suppressed at fitnesses near zero and *f*_max_. This makes the precise value of *f*_max_ less important for determining *α*, and we can approximate **Eq**. (S7) by taking the *f*_max_ → 1 limit. If we then keep the expressions to leading order in *µ*, assuming that *α* ∼ 𝒪(*µ*), **Eq**. (S7) becomes

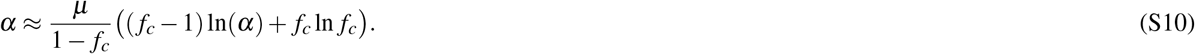

The solution to this equation has the form

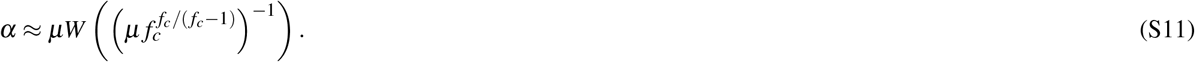

Here *W* (*x*) is the Lambert *W* function, which is the solution *y* of the equation *ye*^*y*^ = *x*. For *x* > 0 (which is the only case that arises in our problem) the function is single-valued (the so-called zero branch of the solution).

While **Eq**. (S11) works in the *f*_max_ → 1 limit, ideally we would like an expression that works for *f*_max_ close to, but not exactly 1, and for the entire range of *µ* ≲ 1 including small *µ*. Since we know 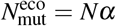 for *μ* ≪ 1 and *f*_max_ < 1 from **Eq**. (S8), we posit the following approximate form for 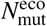 (main text Eq. (4)):

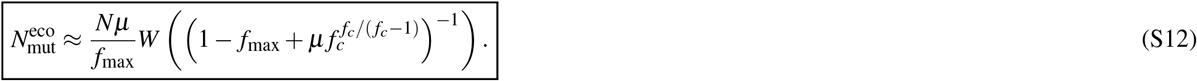

By construction this is consistent with **Eq**. (S11) when *f*_max_ → 1. When 1 − *f*_max_ is small but nonzero and *µ* ≪ 1, we can use the fact that *W* (*x*) diverges like ln *x* for large positive *x* to see that **Eq**. (S12) gives 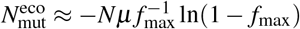 ln(1 − *f*_max_) for *µ* → 0.

This recovers the dominant contribution in Eq. (S8) when 1 − *f*_max_ is small.

Thus in principle **Eq**. (S12) should work for a wide range of *µ* and different values of *f*_*c*_. **Fig. S4** depicts a comparison of **Eq**. (S12) to simulation results, and the analytical approximation is within 10% of the numerical value across five decades of *µ* and both small and large *f*_*c*_ (as shown by the errors shown in the inset). For small *µ* the 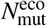 curves start out independent of *f*_*c*_ and proportional to *µ*, as expected from **Eq**. (S8). With increasing *µ*, the curves bend downwards in a way that depends on *f*_*c*_, as the population of mutants becomes a non-negligible fraction of the total.

##### Non-uniform ecological/intrinsic fitness distributions

The derivations above can be easily extended to non-uniform distributions of the ecological and/or intrinsic fitnesses. This modifies the shape of *ρ*_*α*_ (*f*), depending on the specific distributions from which *f*_*i*_ and *f*_*e*_ are drawn for new mutants. In general, imagine *f*_*e*_ is drawn from a distribution *ρ*_0_(*f*_*e*_), like the Gaussian example considered in the main text. In the limit *µ* ≪ 1, when the total mutant fraction *α* is small, **Eq**. (S9) for the distribution becomes (main text Eq. (6)):

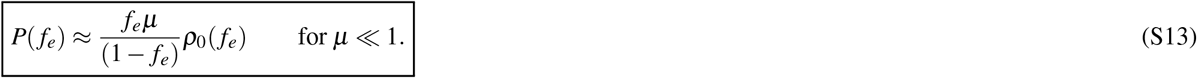

**Figure S4.**
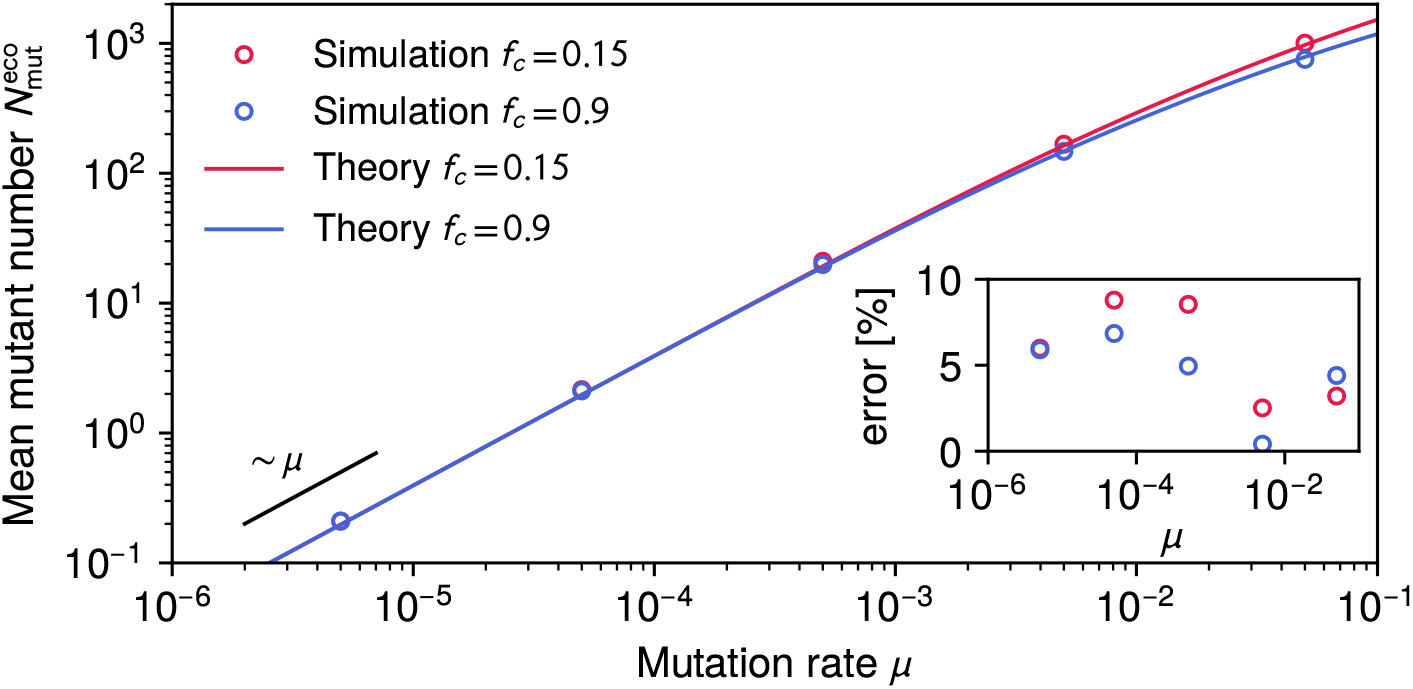
Comparison of the approximate analytical theory, **Eq**. (S12) (curves), and the simulation results (circles) for the mean number of mutants 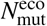 as a function of mutation rate *µ*. We set *f*_max_ = 0.995, *N* = 10^4^, and use two different values for the fitness cost, *f*_*c*_ = 0.15 and 0.9 (red and blue respectively). The inset shows the absolute percent error of the analytical approximation with respect to the numerical values. The black line on the left shows the scaling 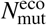 ∝ *µ* expected in the small *µ* regime from **Eq**. (S8).

Other results from the theory can be similarly generalized.

##### No ecological interactions

In the absence of ecological interactions, the fitness function *f* (*α*) = *f*_*i*_ for all *α*. If *f*_*i*_ is drawn from a uniform distribution between 0 and 1− *f*_*c*_, the distribution *ρ*_*α*_ (*f*) becomes uniform: *ρ*_*α*_ (*f*) = 1*/*(1 − *f*_*c*_). An analogous calculation to the one above yields the mean number of mutants in this scenario (main text Eq. (2)):

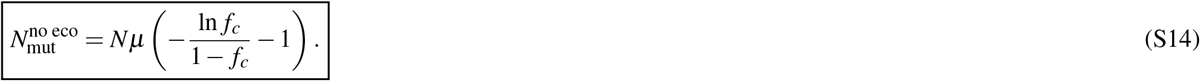

This number is a useful baseline for gauging the relative effectiveness of ecological interactions in enhancing mutant populations.

#### Mean time to extinction of an individual mutant

The final quantity we would like to calculate theoretically is the mean number of generations that an individual mutant survives after first arising. Note that if a mutant is generated by the mutation step, but does not survive the selection step immediately afterwards, we say its lifetime is zero generations. To simplify the calculation, we consider the regime *µ* ≪ 1, where the chance that a mutant disappears via a second mutation is negligible. And by the definition of the model, the same type of mutant cannot be generated again from either the ancestor or other mutant populations. So in this regime the mutant persists with a randomly fluctuating population until in one of the selection steps none of its population is chosen to survive to the next generation.

Let us focus on a single mutant type with fitness *f*. If there are *ℓ* such mutants in the current generation, the probability *W*_*kℓ*_ that there will be *k* mutants of this type in the next generation is given by the binomial distribution characteristic of Wright-Fisher dynamics,

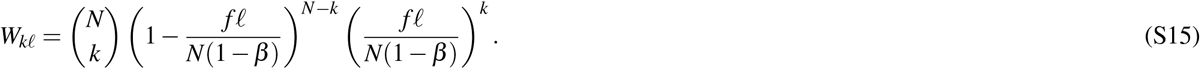

For *µ* ≪ 1 we can assume that *β* ≪ 1, *f* ≈ *f*_*e*_ and that *N* ≫ *k* for any *k* that has a non-negligible probability, since the number of mutants of a single type at any given time will be a tiny fraction of the total. We can then approximate **Eq**. (S15) as

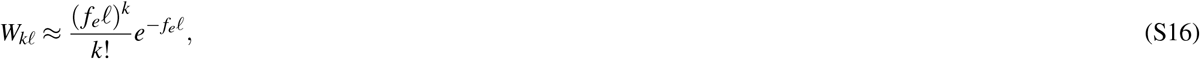

which is just the limit in which the binomial distribution looks like a Poisson distribution. The probabilities *W*_*kℓ*_ can be interpreted as components of an *N* × *N* transition matrix *W*. Since the vast majority of this matrix will consist of probabilities exponentially close to zero, we can focus on the states *k, ℓ* = 0, …, *M* for some *M* ≪ *N*. Thus we will consider *W* instead to be an (*M* + 1) × (*M* + 1) matrix, choosing *M* large enough to get a satisfactory approximation to the non-truncated system. This reduces the problem to an (*M* + 1) state discrete time Markov process.

Note that *W*_*k*0_ = 0, so *ℓ* = 0 (extinction) is an absorbing state. From the theory of phase-type distributions^58^, the mean number of generations to extinction can be calculated using the *M* × *M* submatrix *S* of the tranpose *W*^*T*^, defined via *S*_*kℓ*_ = *W*_*ℓk*_ for *k, ℓ* = 1, …, *M*. Starting from a population of 1 (after the mutation step) at generation zero, the mean time to extinction is given by

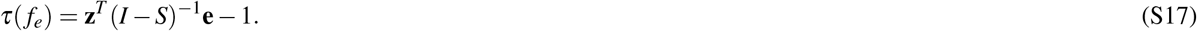

Here *I* is the *M* × *M* identity matrix, **z** is an *M*-dimensional vector with a 1 in the first element and zero elsewhere, and **e** is an *M*-dimensional vector with 1 for all its elements. The − 1 at the end of **Eq**. (S17) is due to the counting convention where extinction during the first selection step is considered to be an extinction time of zero.

For a given choice of *M*, it turns out the matrix inverse in **Eq**. (S17) can be calculated analytically. Even though the resulting expression becomes unwieldy for large *M*, it can always be Taylor expanded around *f*_*e*_ = 0 to give relatively simple results. The Taylor coefficient of order 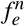 in the expansion remains unchanged for any choice of *M* ≥ *n*. Thus we can find the Taylor expansion of *τ*(*f*_*e*_) in the non-truncated system up to any chosen order *M*, simply by Taylor expanding **Eq**. (S17).

The first few terms of this Taylor expansion are:

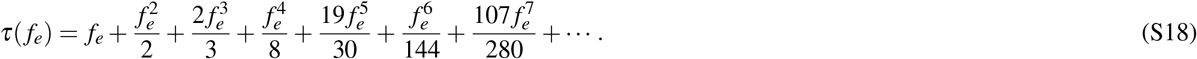

While the lowest terms are sufficient to describe the mean extinction time of mutants with *f*_*e*_ ≪ 1, progressively more terms are required to approximate *τ*(*f*_*e*_) as *f*_*e*_ approaches 1 from below. In fact, technically in this approximation the series *τ*(*f*_*e*_) diverges at *f*_*e*_ = 1, since we have effectively taken *N* → ∞ in **Eq**. (S16). In practice this is not a problem since we only consider mutants with fitnesses up to *f*_max_ < 1. Rather than working with the Taylor expansion directly, we constructed an analytical approximation designed to agree with the expansion through order 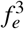, and capture the divergence at large *f*_*e*_ (main text Eq. (1)):

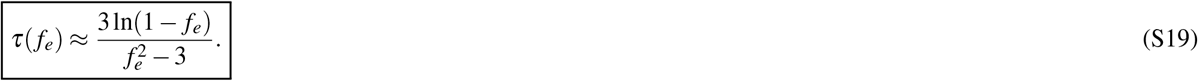

Despite its simple form, this approximation agrees with the simulation results for *µ* ≪ 1 across the whole range of *f*_*e*_ with a typical error of 5%.

